# Re-analysis of lipidomic data reveals Specialised Pro-Resolution Lipid Mediators (SPMs) to be lower than quantifiable limits of assay in a human model of resolving inflammation

**DOI:** 10.1101/2023.03.06.530669

**Authors:** Natalie ZM Homer, Ruth Andrew, Derek W Gilroy

**Affiliations:** Mass Spectrometry Core, Edinburgh Clinical Research Facility, University/BHF Centre for Cardiovascular Sciences, Queen’s Medical Research Institute, 47 Little France Crescent, Edinburgh EH16 4TJ, United Kingdom; University/BHF Centre for Cardiovascular Science, Queen’s Medical Research Institute, University of Edinburgh, 47 Little France Crescent Edinburgh, EH16 4TJ; Dept. of Experimental & Translational Medicine, The Rayne Building, University College London, 5 University Street, London WC1E 6JF

## Abstract

Using a model of UV-killed *E. coli* driven dermal inflammation in healthy human volunteers we originally reported that following inflammatory resolution there was the infiltration of macrophages, which, through prostanoids including prostaglandin (PG)E_2_, imprints long-term tissue immunity. In addition to the prostanoids, we also presented data on levels of <u>S</u>pecialised <u>P</u>ro-Resolution Lipid <u>M</u>ediators (SPMs) throughout inflammatory onset, resolution and post-resolution phases of this model. However, our collaborators who carried out the lipidomic analysis received a complaint concerning how they generally present SPM data in their publications, namely their use of graphical illustrations to depict data. Importantly, such lipidomic illustrations were used in our human UV-killed *E. coli* study. Therefore, in the interest of transparency and to replace these illustrations with more meaningful images, the original data from our human UV-killed *E. coli* model were re-analysed by two independent experts. It transpires that the integrated areas of the chromatographic peaks of the SPM lipid mediators were below the amounts that could be reliably either detected and/or quantified using community standards for quantitation. Here we show the outcome of this reanalysis. Importantly, with prostanoids including PGE_2_ being robustly detected, this re-analysis does not alter the original report that post-resolution PGs imprint tissue immunity.

## INTRODUCTION

We originally examined the role of lipid mediators in the resolution and post-resolution phases of an experimental model of self-limiting inflammation in healthy human volunteers triggered by an intradermal of UV-killed E Coli (UV-KEc)(1). This necessitated the use of mass spectrometry (MS) as the gold standard technique with which to determine the presence and quantities of lipid mediators in inflammatory exudates that were extracted from the inflamed skin. These lipids included <u>S</u>pecialised <u>P</u>ro-Resolution Lipid <u>M</u>ediators (SPMs), prostaglandins (PGs) and leukotrienes (LTs). Sample processing and data analysis was performed by Professor Jesmond Dalli, Lipid Mediator Unit Director, William Harvey Research Institute, Barts and The London School of Medicine and Dentistry, Queen Mary University of London (QMUL), Charterhouse Square, London. UK. EC1M 6BQ.

However, Professor Dalli recently informed the lead author, Professor Derek W Gilroy, of an external complaint received by QMUL regarding an illustration used to depict lipid molecules identified in samples displayed in **FIGURE S1** of the above paper, with concentrations and temporal profiles of a representation of these lipids presented in **Figure 4A-H**. The concern was that such illustrations in **FIGURE S1** might be misunderstood as reporting raw data.

**Figure 1.**
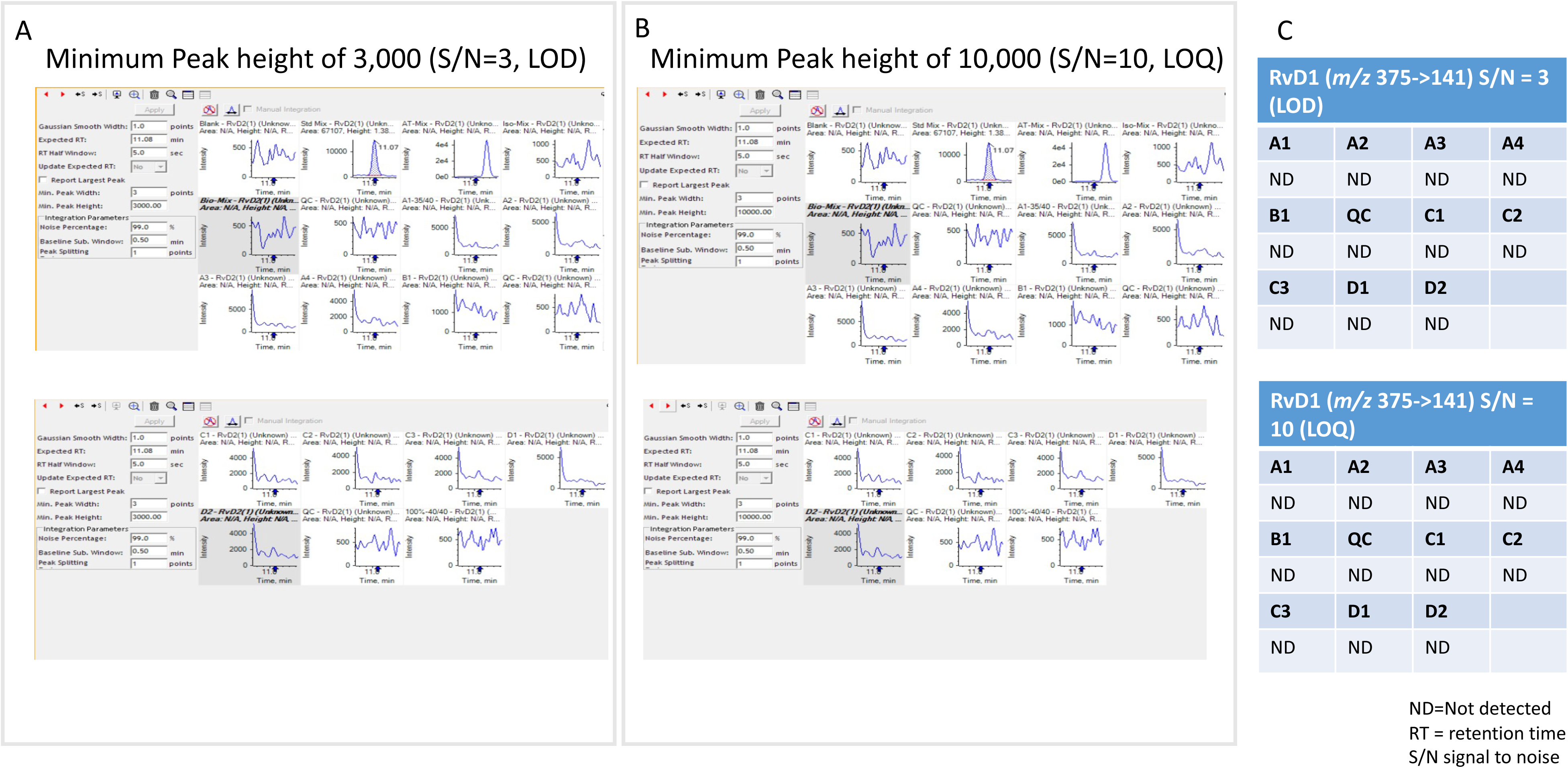
RvD1 (*m/z* 375.2 → 141 RT = 11.08 minutes)

**Figure 2.**
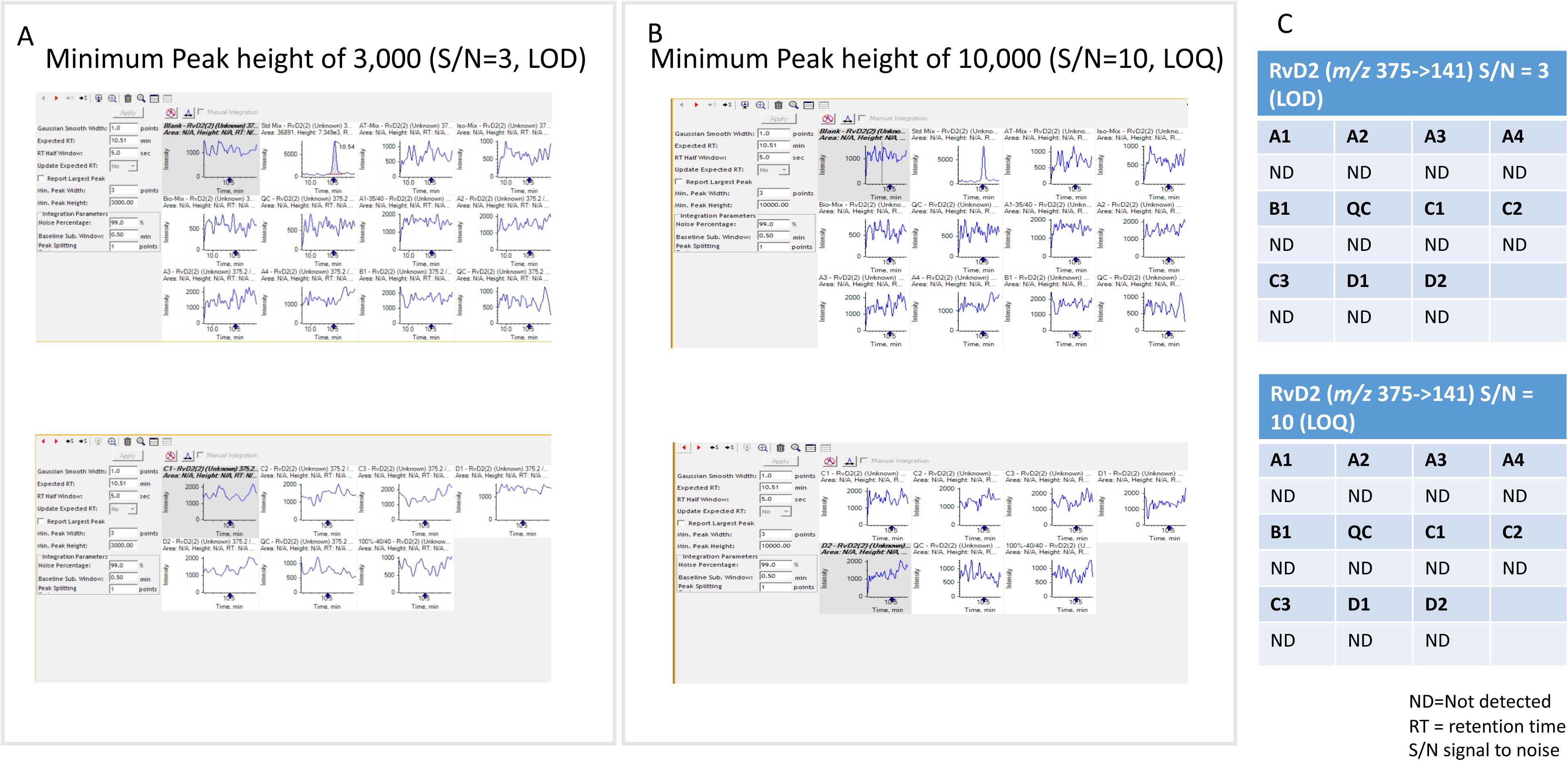
RvD2 (*m/z* 375.2 → 141.1 RT = 10.51 minutes)

**Figure 3.**
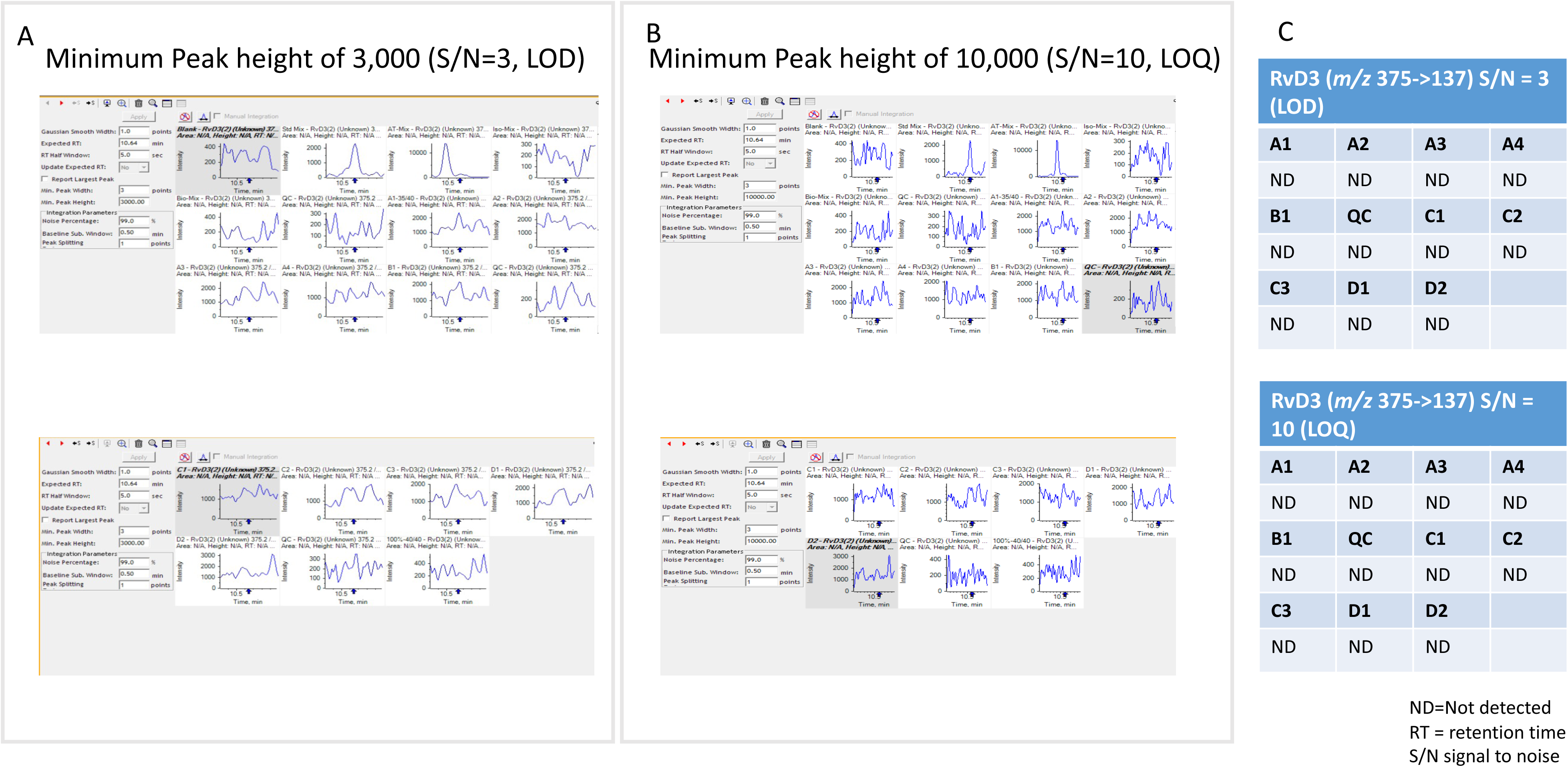
RvD3 (*m/z* 375.2 → 137 RT = 10.64 minutes)

**Figure 4.**
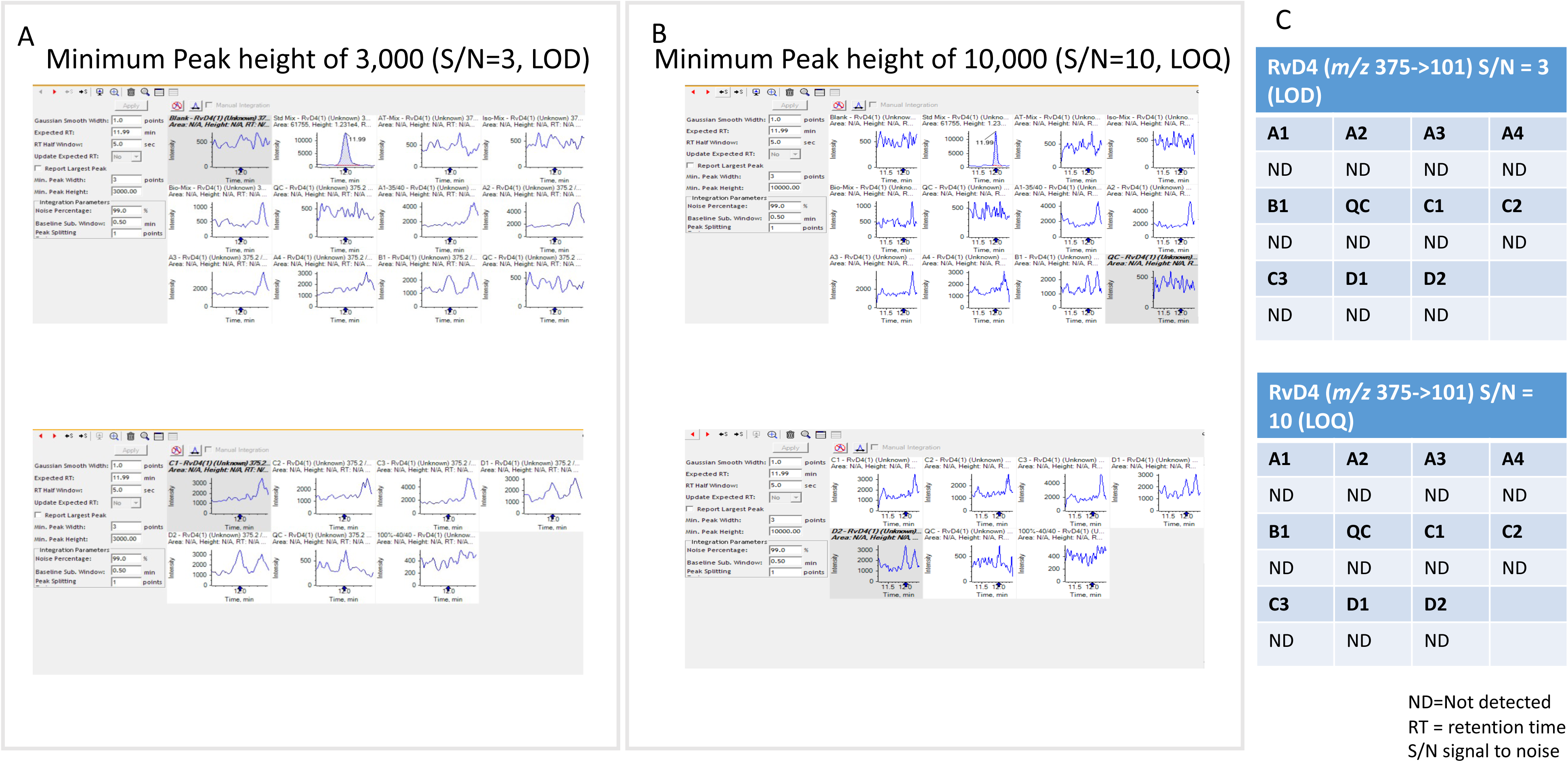
RvD4 (*m/z* 375 → 101 RT = 11.99 minutes)

In response, Professor Gilroy sought the expertise of two external specialists in the field of MS to carry out an unbiased re-analysis of raw data from a subset of samples, in accordance with harmonised ICH guidelines(2, 3). The aim was to replace the “illustration” with data presented in a manner that can be critiqued by other MS experts and understood by non-MS experts who have an interest in inflammation, resolution and lipid biology. Peak areas of a calibration curve were provided in addition to the raw data.

A subset of the raw data (detailed in Results) was provided and, upon reanalysis, we found that the integrated areas of the chromatographic peaks of many of the SPM lipid mediators previously represented in the Supplementary file and Figure 4 were below the amounts that could be reliably either detected and/or quantified using signal to noise ratios of 3:1 and 10:1 respectively as indicative cut-offs; these settings reflect community standards for quantitation(2, 3).

## METHOD OF DATA RE-ANALYSIS

Originally, samples were collected from nine time points following bacterial injection including naïve skin. While **Figure 4** of the original paper presented data on prostanoids (PGE_2_, PGD_2_, PGF_2α_ and TxB_2_,) and limited numbers of SPMs (LxB4, RvD3 and RvD5), data in **supplementary Figure S1A** comprised illustrations of additional members of the SPM family, namely docosahexaenoic acid-derived resolvins, protectins and maresins; eicosapentaenoic acid-derived resolvins as well as prostanoids and leukotriene.

Here, we re-assessed a representative number of the SPM family, including those in **Figure 4** and in the **Supplementary Figure S1A** namely RvD1; RvD2; RvD3; RvD4; RvD5; RvD6; 17-RvD1; 17-RvD3; PD1; 10S, 17S-diHDHA; MaR1; 7S, 14S-diHDHA; LXB4; RvE3, 5S,15S-diHETE; RvE1; RvE2; LxA4; 15-epi-LxA4 in the subset of the data provided. These were from five time points including 8h and 14h (early inflammation onset), 48h (resolution) and day 14 and day 17 post UV-KEc injection of the original analysis. We also re-analysed data for PGE2, PGD2, PGF2α and TxB2 as well as LTB4 for comparison purposes.

Retention times and mass transitions of lipid mediators in this LC-MS/MS method were provided by Professor Dalli. MultiQuant 3.0.3 software (Sciex, Warrington, UK) was used to create an automatic method of integration according to the precursor (Q1) and product (Q3) mass transitions (denoted Transition label) and retention times outlined in **Table 1**. A 5 second retention time threshold was defined to aid in discrimination of isobaric lipids.

**Table 1.** Lipid mediator LC-MS/MS method details.

| Analyte name | Retention time | Label of Transition | Q1 | Q3 |
| --- | --- | --- | --- | --- |
| RvD1 | 11.08 | RvD2(2) | 375.2 | 141.1 |
| RvD2 | 10.51 | RvD2(2) | 375.2 | 141.1 |
| RvD3 | 10.64 | RvD3(2) | 375.2 | 137.0 |
| RvD4 | 11.99 | RvD4(1) | 375.2 | 101.0 |
| RvD5 | 13.20 | RvD5(1) | 359.2 | 199.2 |
| RvD6 | 13.70 | RvD6 | 359.2 | 159.1 |
| 17-RvD1 | 11.25 | RvD1(2) | 375.2 | 215.2 |
| 17-RvD3 | 10.47 | RvD3(2) | 375.2 | 137.0 |
| PD1 | 13.30 | PD1(1) | 359.2 | 153.1 |
| 10S,17S-diHDHA | 13.10 | PD1(2) | 359.2 | 181.1 |
| MaR1 | 13.40 | MaR1(3) | 359.2 | 141.1 |
| 7S,14S-diHDHA | 13.60 | MaR1(3) | 359.2 | 141.1 |
| LXB<sub>4</sub> | 10.59 | LXB4 | 351.1 | 221.1 |
| RvE3 | 13.40 | RvE3 | 353.3 | 201.1 |
| 5S,15S-diHETE | 13.30 | 5S,15S-diHETE (2) | 335.3 | 235.1 |
| RvE1 | 8.15 | RvE1 (2) | 349.2 | 161.2 |
| RvE2 | 11.9 | RvE2(2) | 333.3 | 159.2 |
| LXA<sub>4</sub> | 11.1 | LXA4(2) | 351.1 | 115.1 |
| 15-epi-LXA<sub>4</sub> | 11.36 | LXA4 (2) | 351.1 | 115.1 |
| PGE<sub>2</sub> | 10.39 | PGE2 | 351.1 | 189.1 |
| PGD<sub>2</sub> | 10.50 | PGD2 | 351.1 | 189.1 |
| PGF2α | 10.78 | PGF2α | 353.3 | 193.1 |
| T<sub>x</sub>B<sub>2</sub> | 9.91 | TXB2 | 369.3 | 169.1 |
| LTB<sub>4</sub> | 13.6 | LTB4 | 335.3 | 195.1 |

We used the widely accepted rule of a minimum signal-to-noise ratio (SNR) of three to assign the limit of detection and the second rule of SNR of 10 to assign the limit of quantitation. Baseline was assigned adjacent to, and ahead of the peak of interest in the biological samples. The baseline was generously assigned as 1,000 peak height broadly across the dataset for all analytes of interest and thus a 3,000 or 10,000 peak height were inferred to assess limit of detection (LOD) and the limit of quantitation (LOQ), respectively (2, 3). Data from a representative standard curve (but not run contemporaneously) was provided and accuracy around the relevant range of calibration points for values deemed quantifiable was cross-referenced.

## RESULTS

Data in **Figures 1-19** display the outputs of reanalysis of the 19 SPMs. In each case, setting a limit of a minimum peak height of 3,000 (LOD; SNR> 3) is displayed as **Panel A**. Data were reanalysed with a limit of peak height of 10,000 (LOQ; SNR>10) as **Panel B**. A summary table of the peak areas of the peaks detected is given as **Panel C**. Raw screenshots of the data and the parameters set are visible to avoid any uncertainty or concerns of data manipulation in the display.

**Figure 5.**
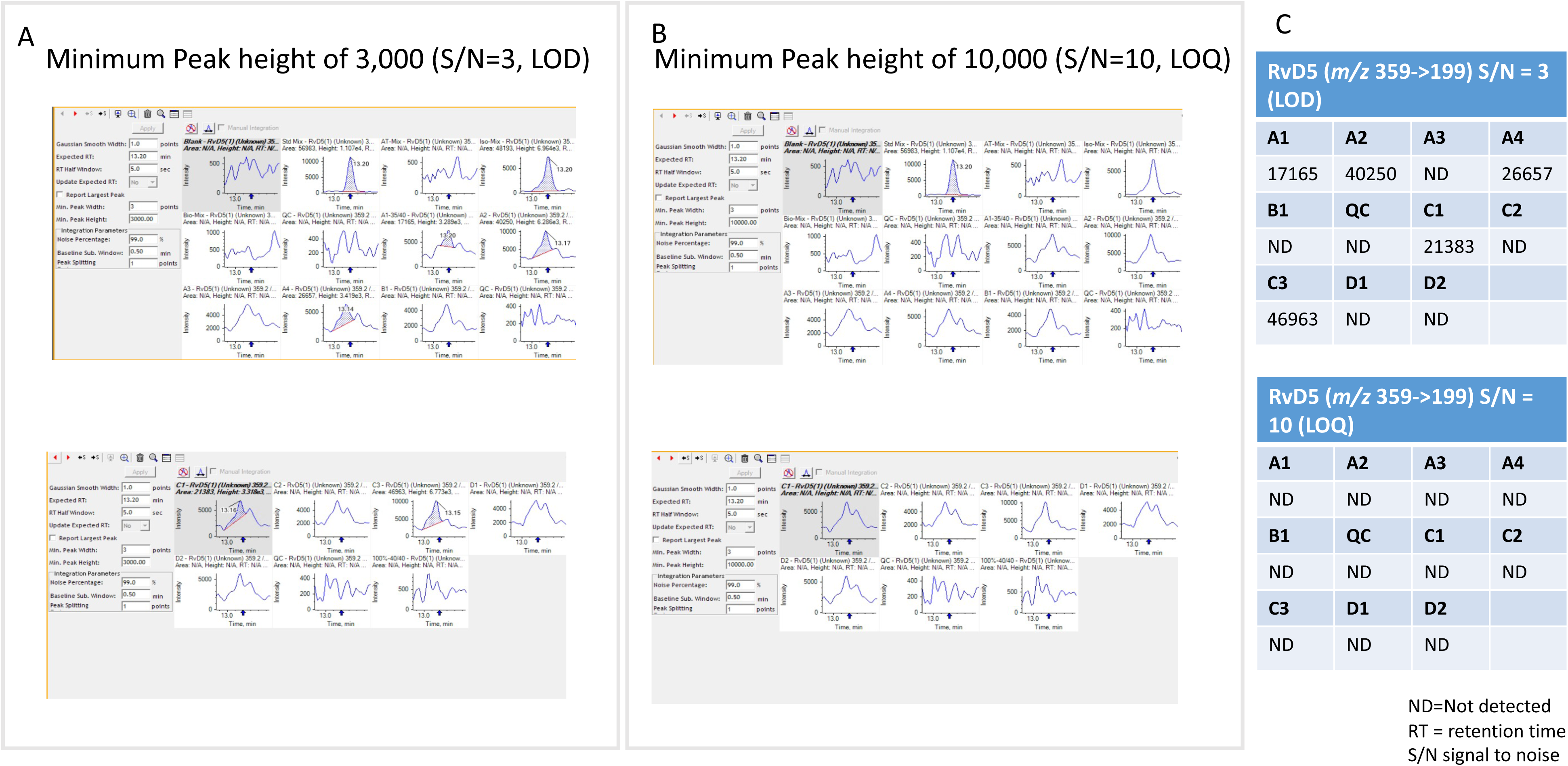
RvD5 (*m/z* 359.2 → 199.2 RT = 13.20 minutes)

**Figure 6.**
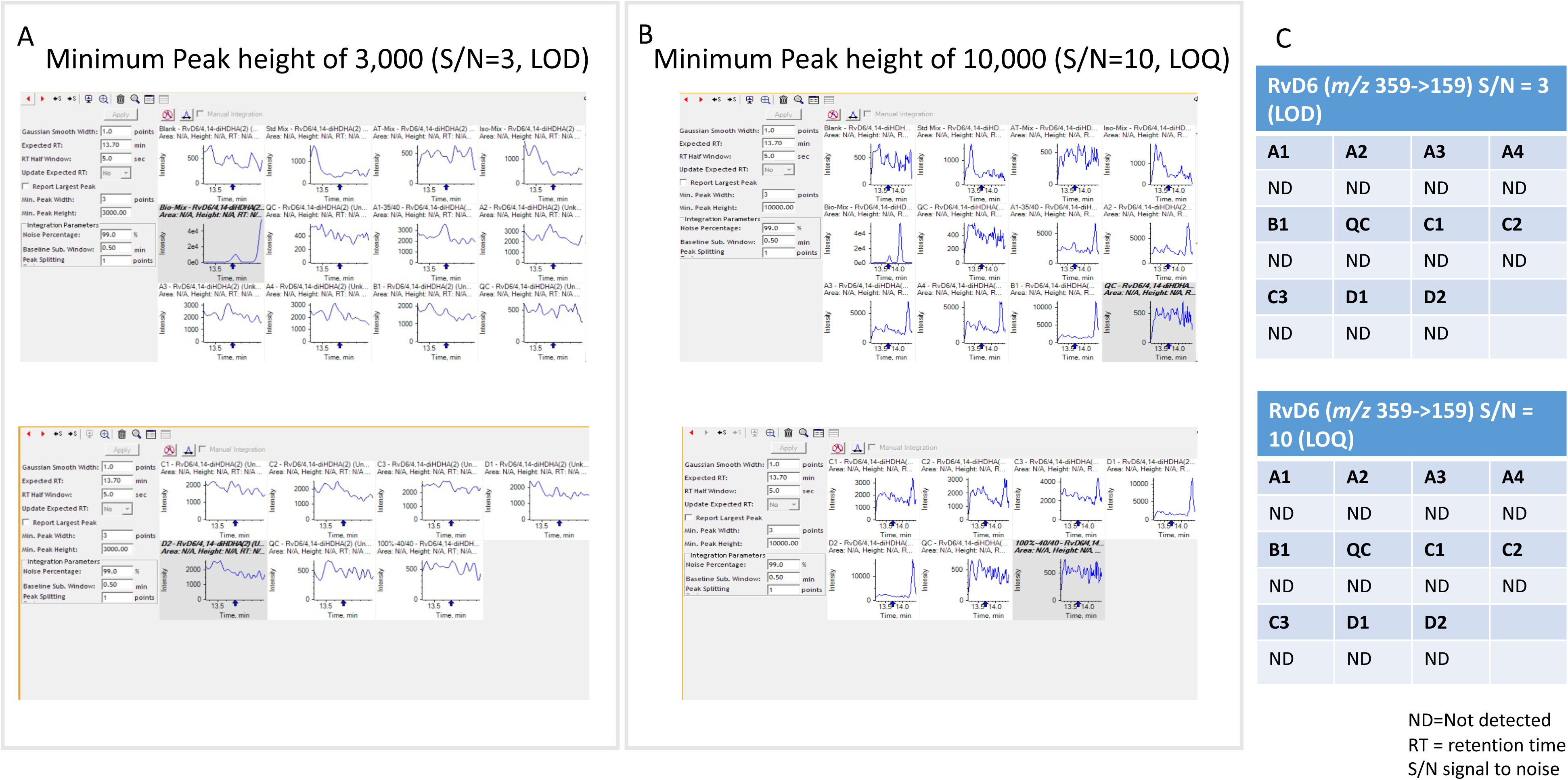
RvD6 (*m/z* 359.3 → 159.1 RT = 13.70 minutes)

**Figure 7.**
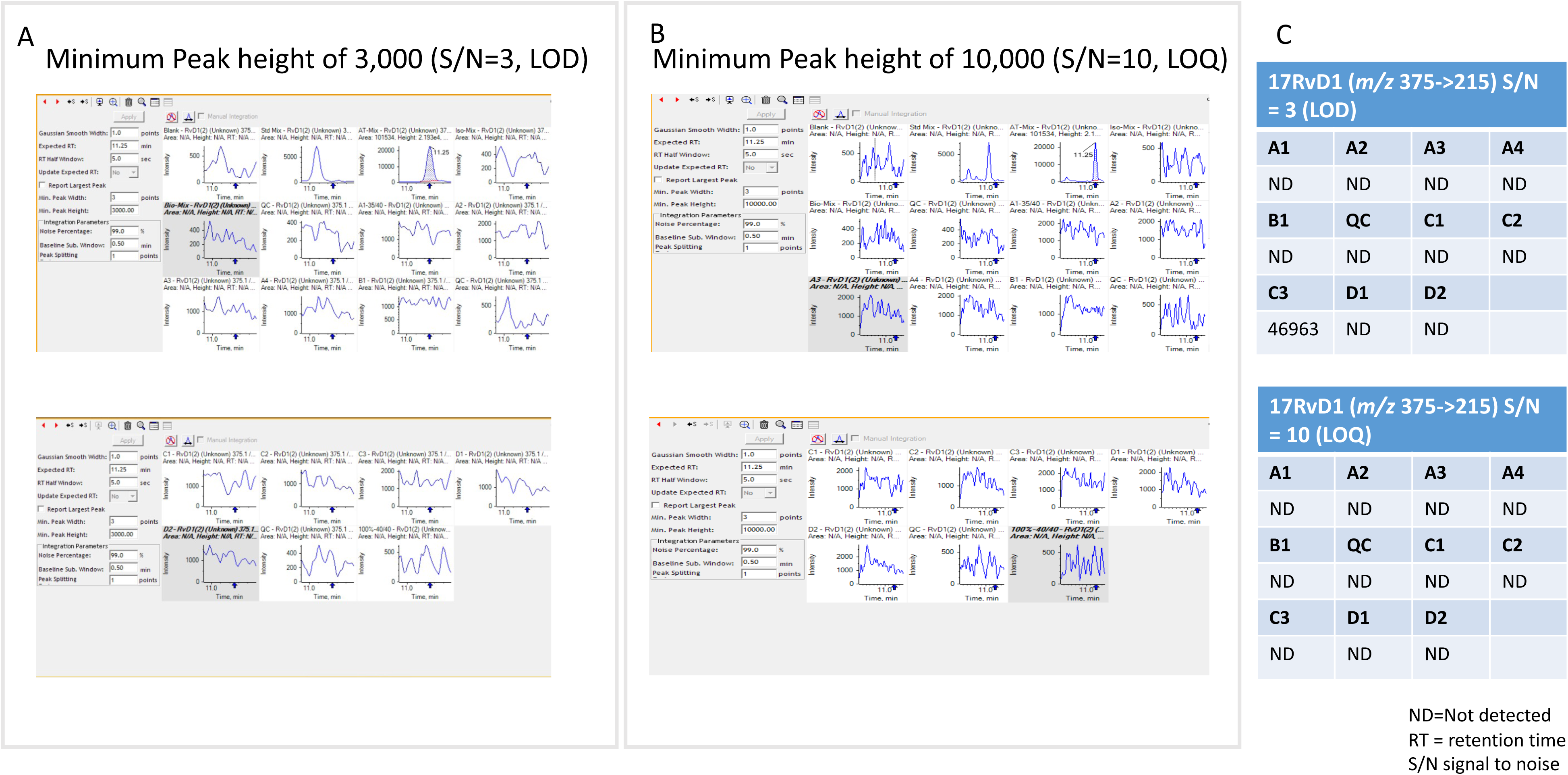
17-RvD1 (*m/z* 375.2 → 215.2 RT = 11.25 minutes)

**Figure 8.**
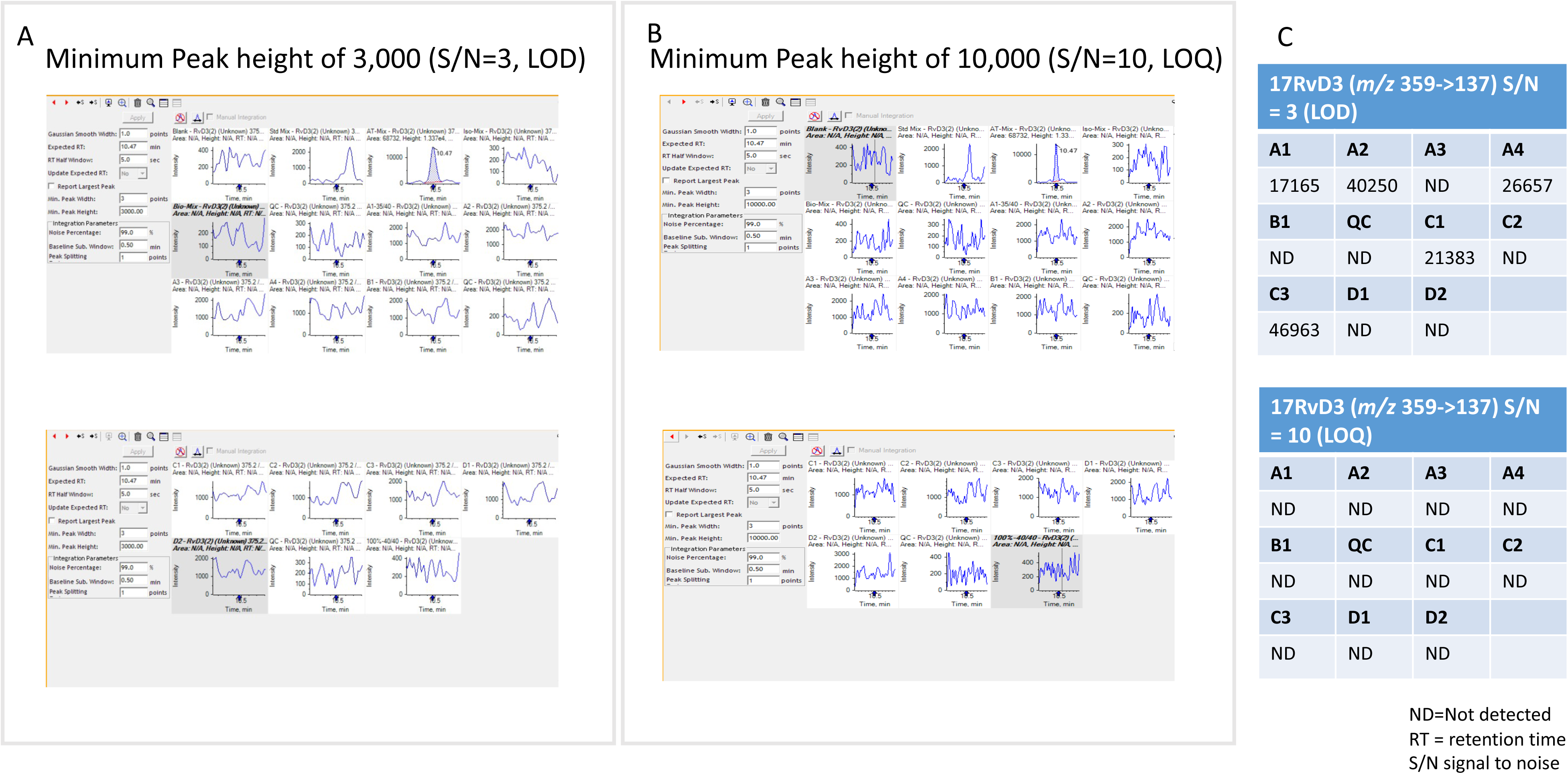
17-RvD3 (*m/z* 375.2 → 137 RT = 10.47 minutes)

**Figure 9.**
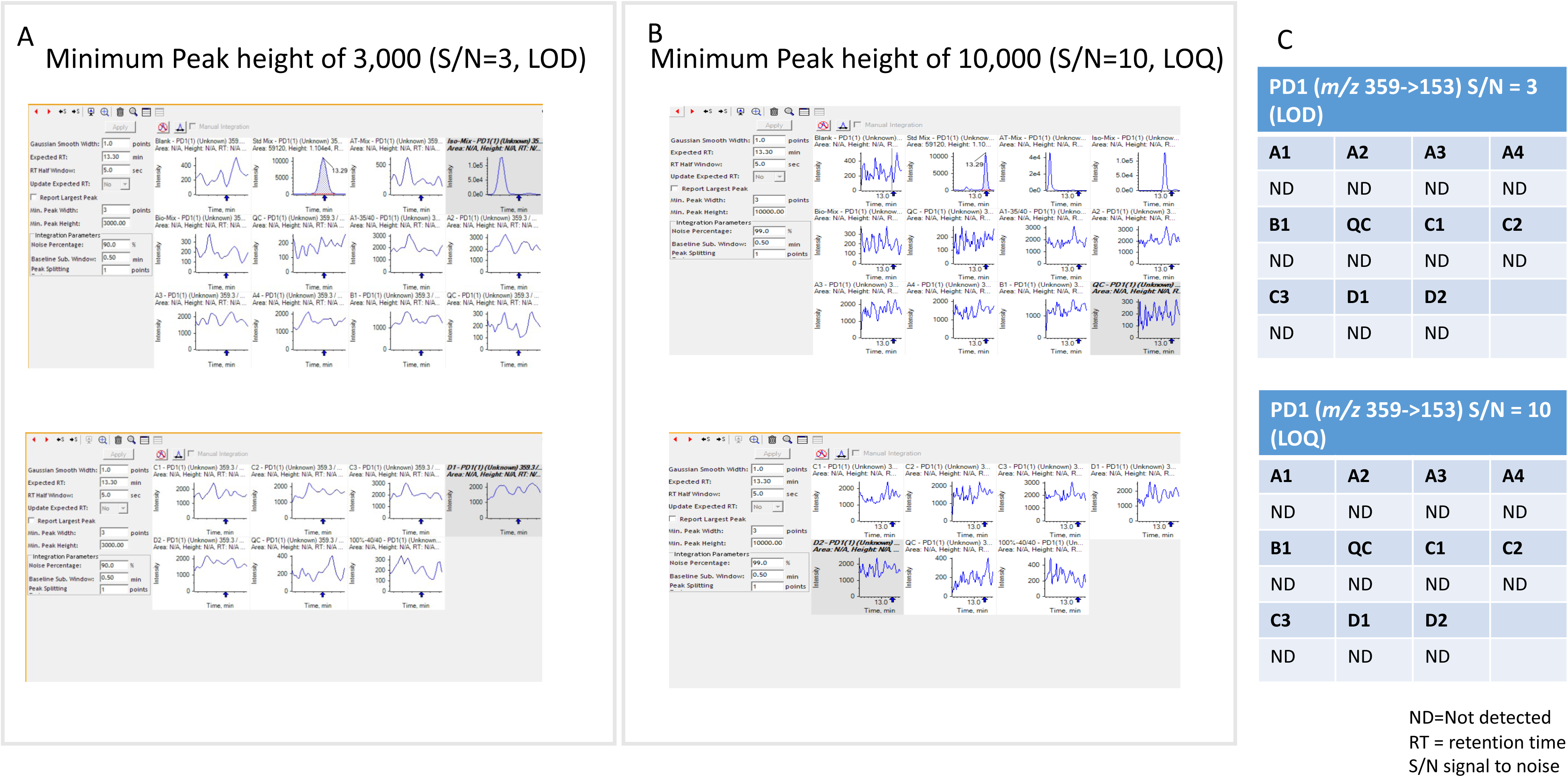
PD1 (*m/z* 359.2 → 153.1 RT = 13.30 minutes)

**Figure 10.**
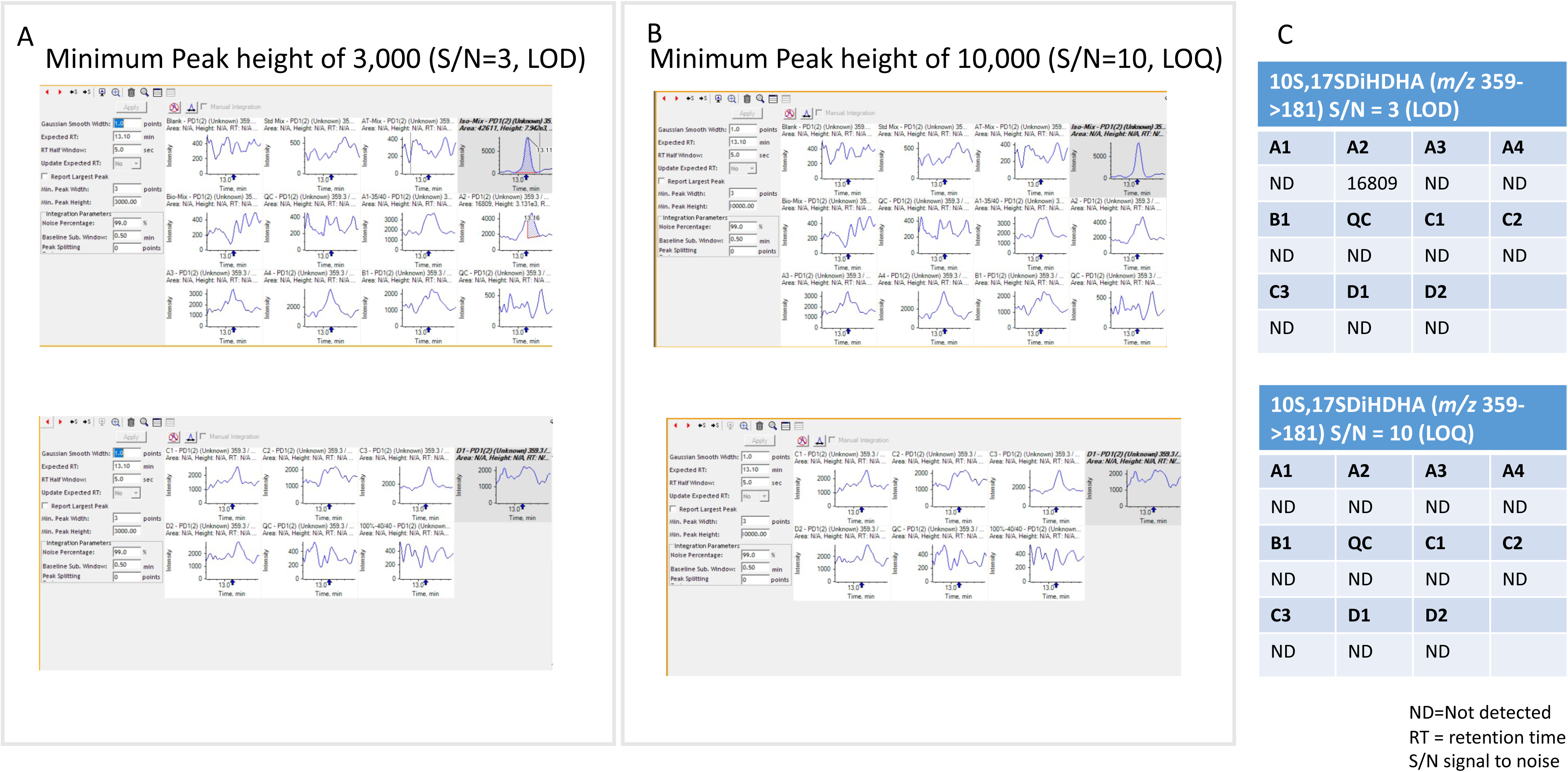
10S, 17S diHDHA (*m/z* 359.2 → 181.1 RT = 13.10 minutes)

**Figure 11.**
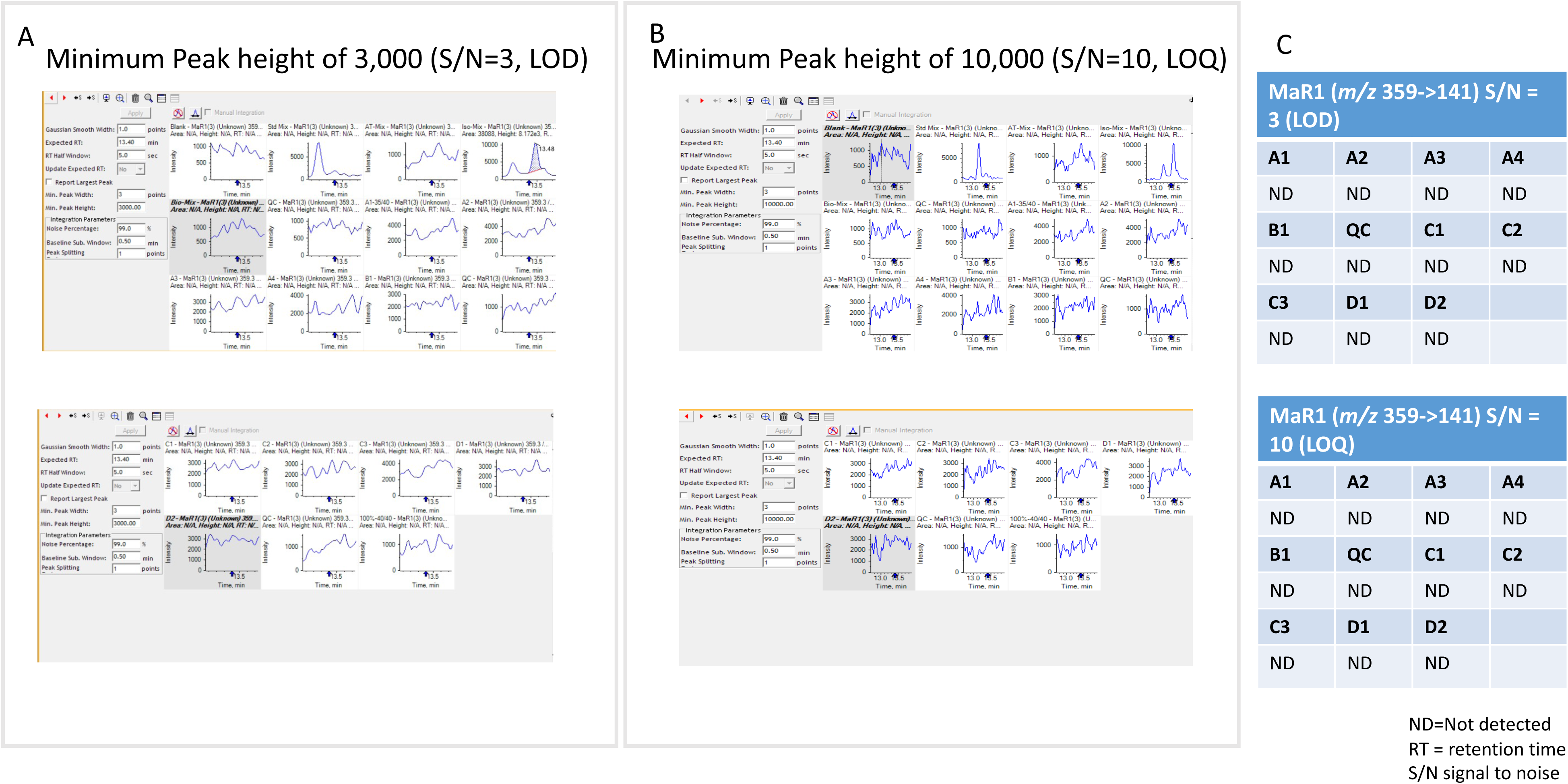
MaR1 (*m/z* 359.2 → 141.1 RT = 13.40 minutes)

**Figure 12.**
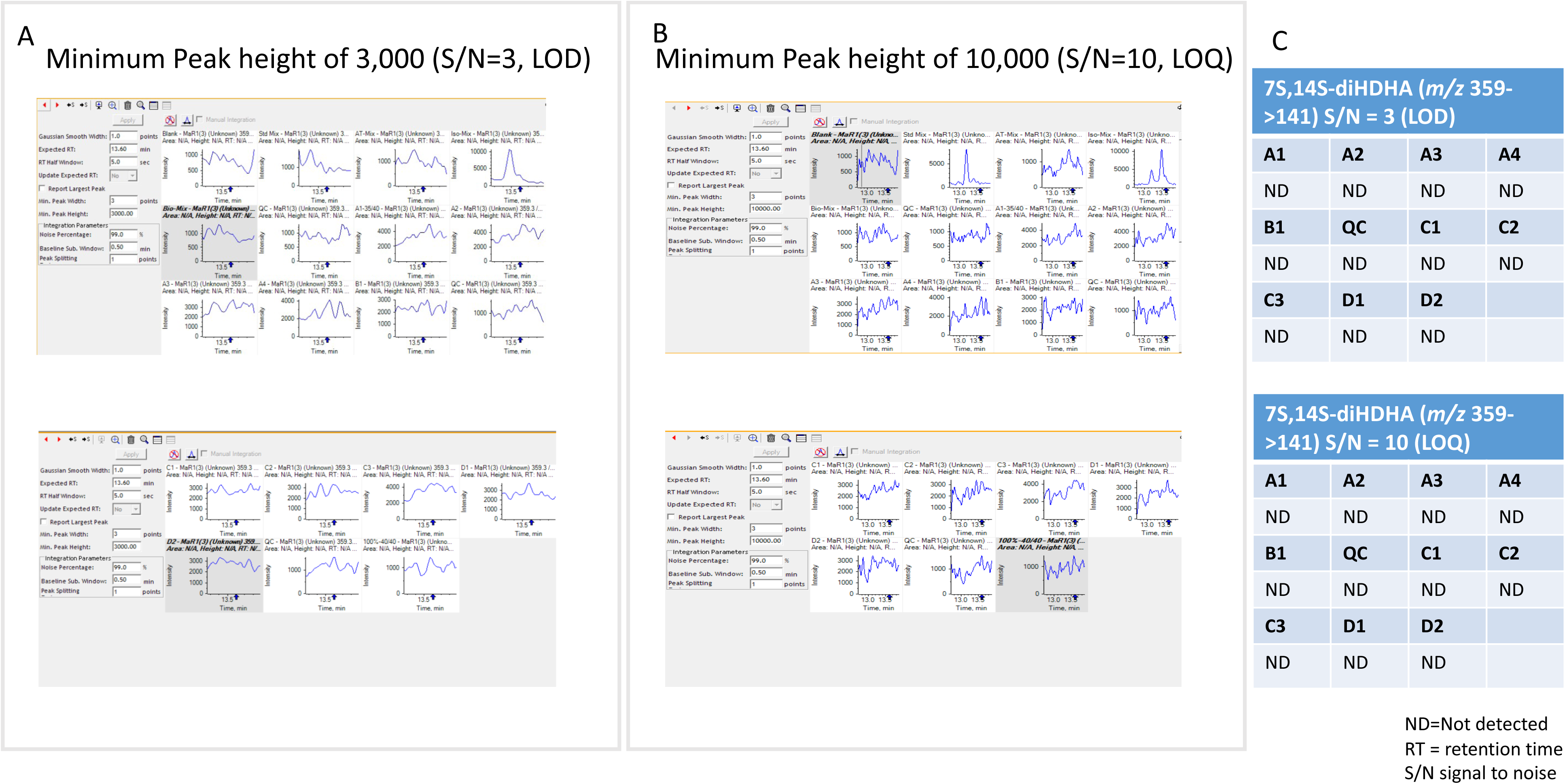
7S,14S-diHDHA (*m/z* 359.2 → 141.1 RT = 13.60 minutes)

**Figure 13.**
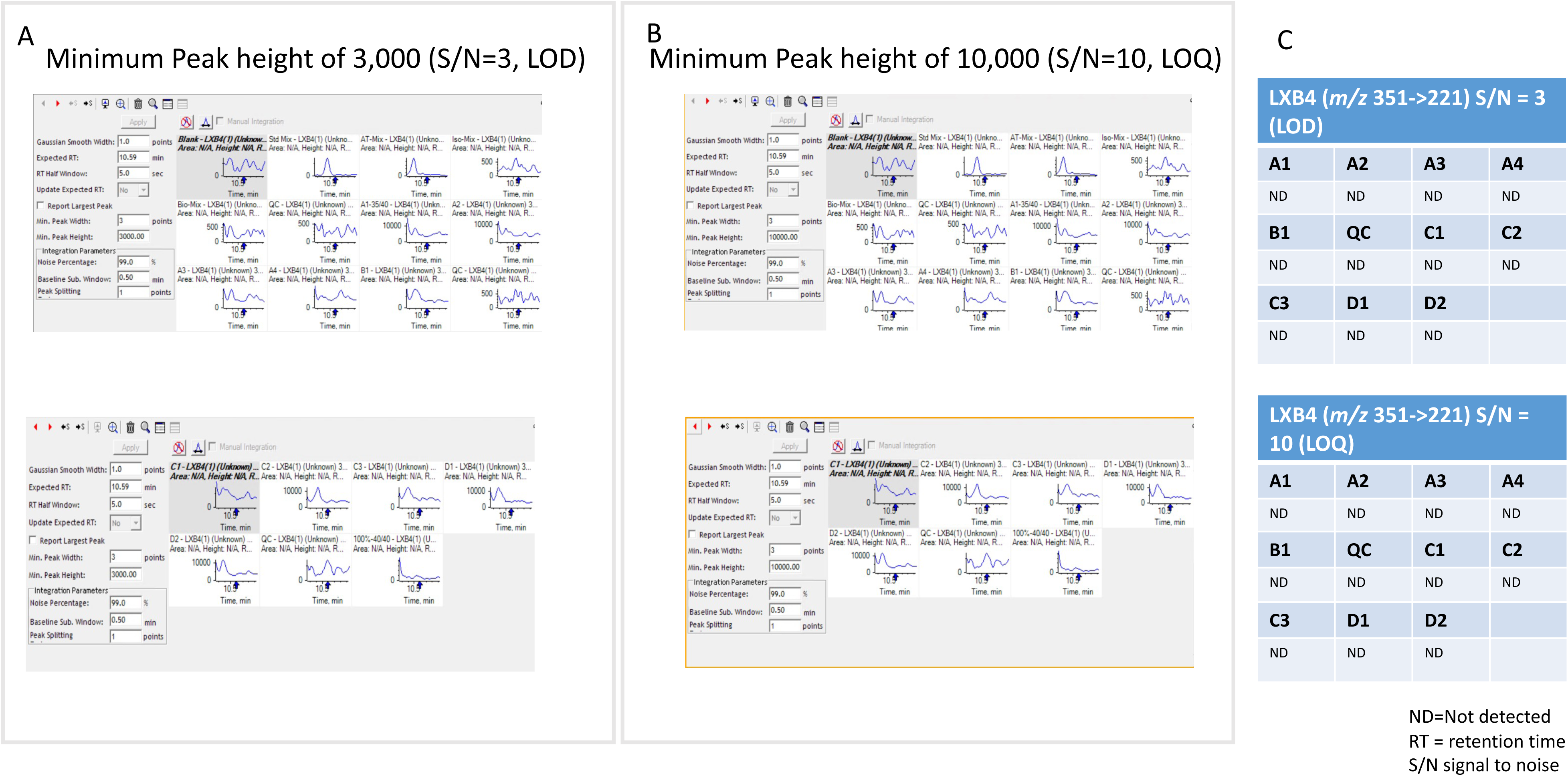
LXB_4_ (*m/z* 351.1 → 221.1 RT = 10.59 minutes)

**Figure 14.**
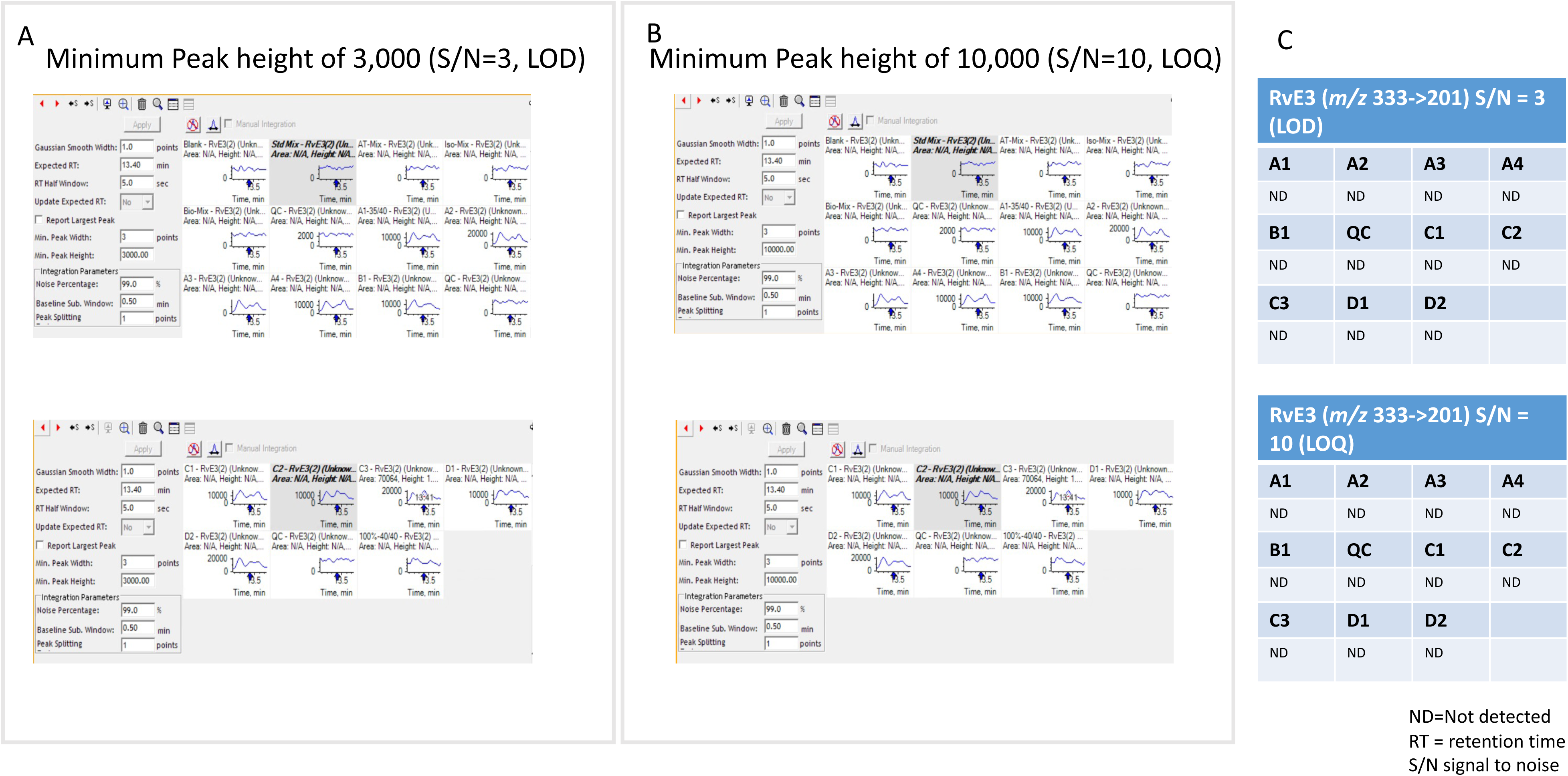
RvE3 (*m/z* 333.3 → 251.3 RT = 13.4 minutes)

**Figure 15.**
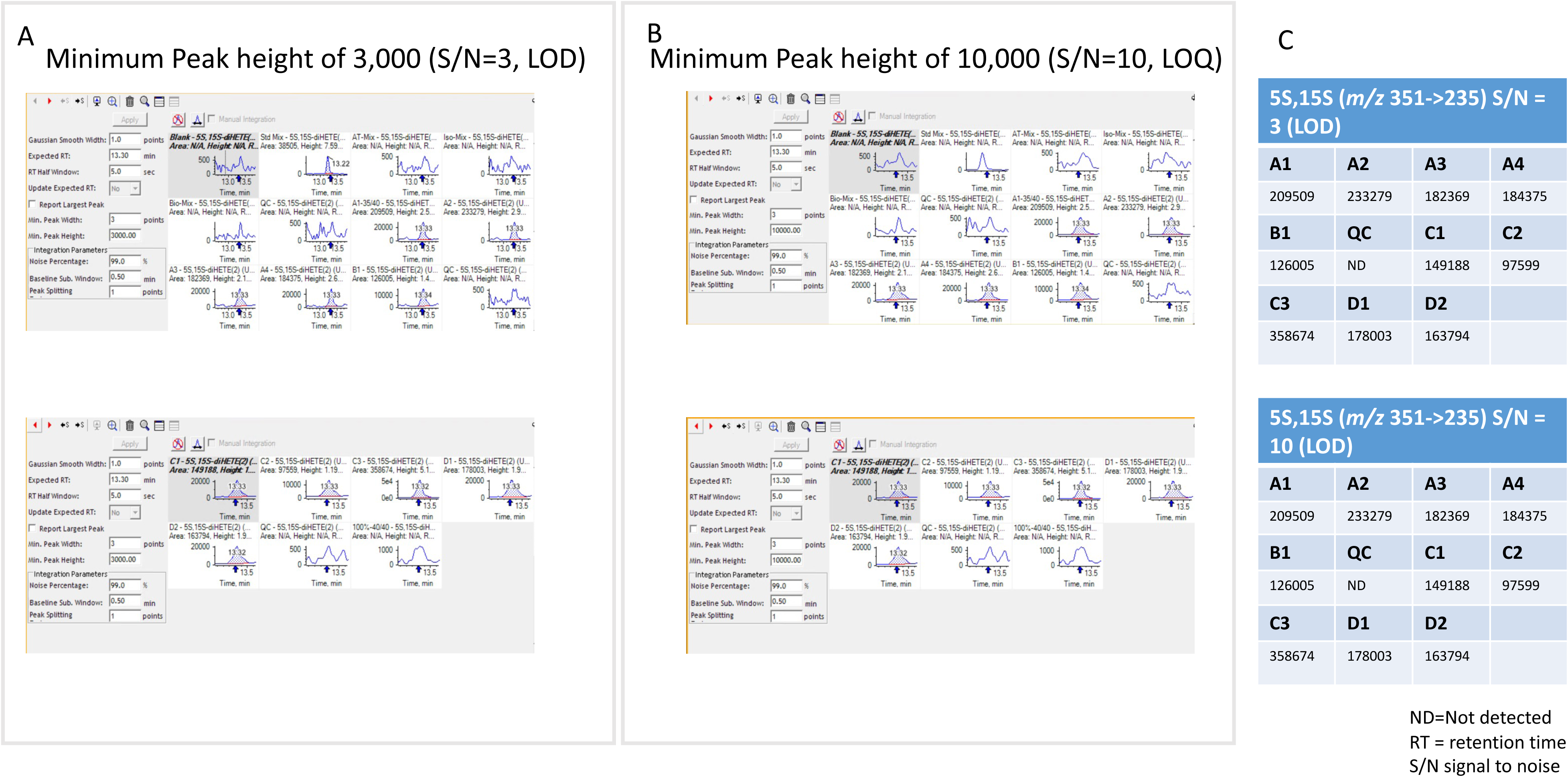
5S,15S-diHETE (*m/z* 335.3 → 235.1 RT = 13.3 minutes)

**Figure 16.**
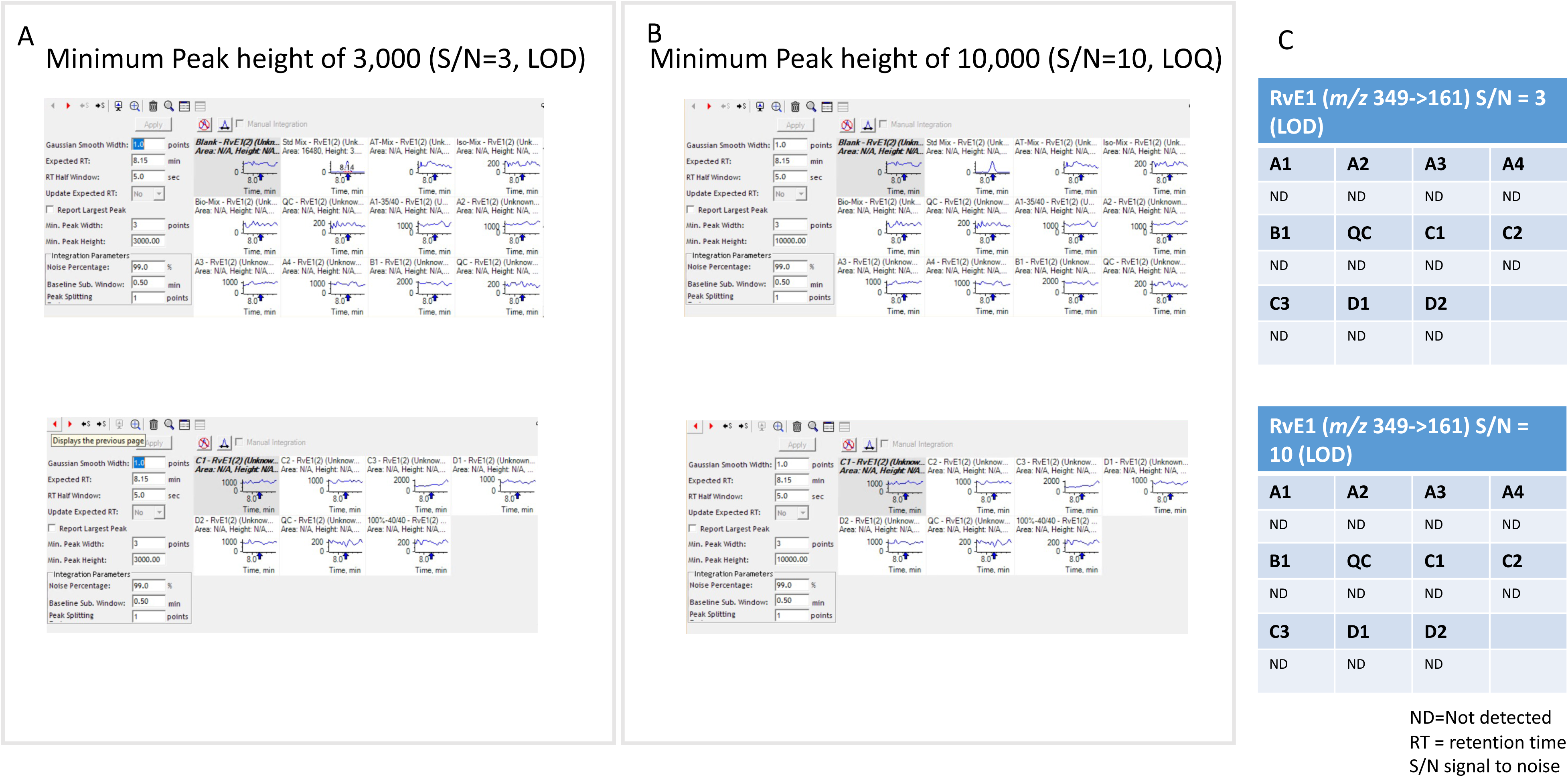
RvE1 (*m/z* 349.2 → 161.2 RT = 8.15 minutes)

**Figure 17.**
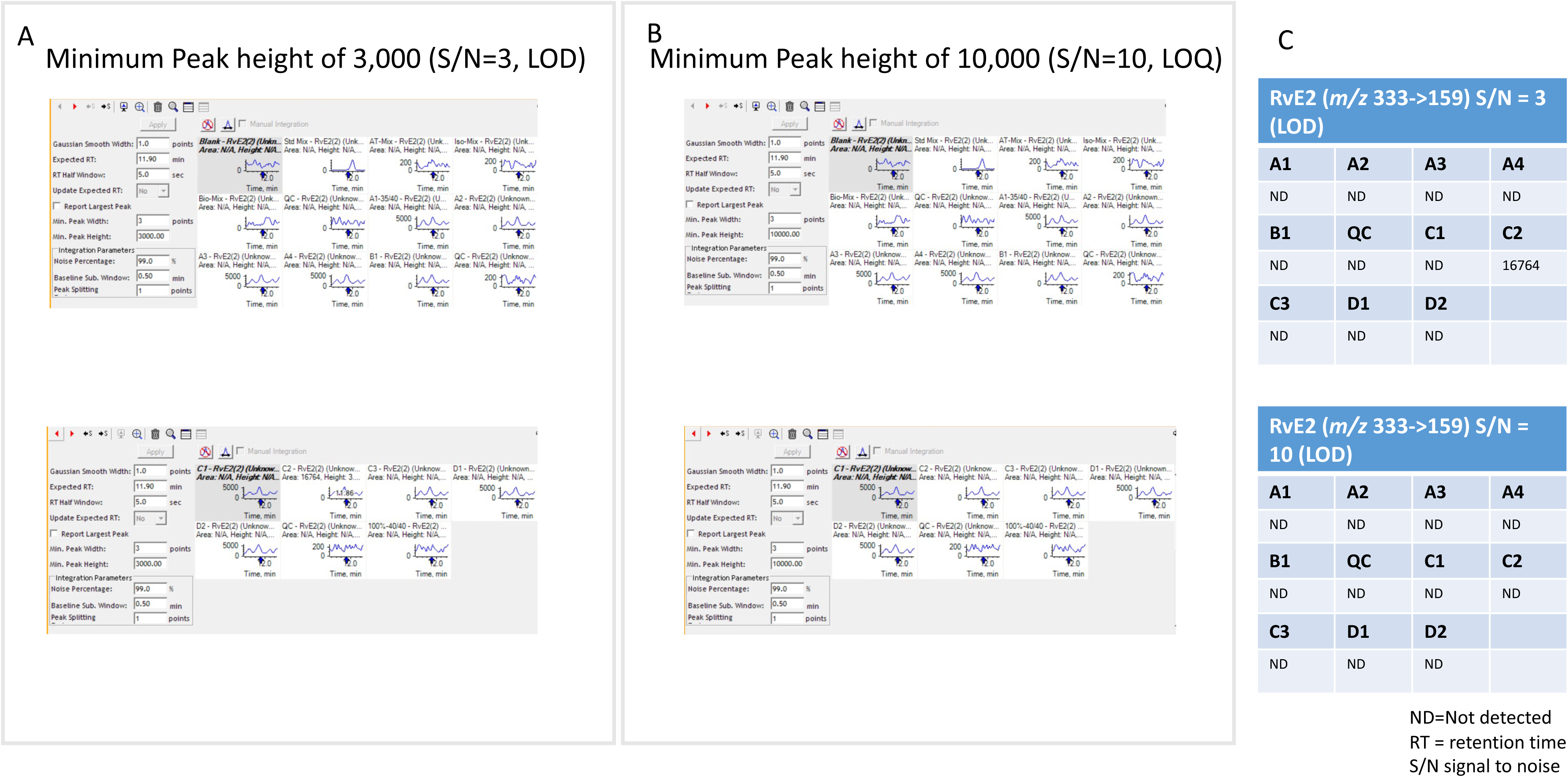
RvE2 (*m/z* 333.3 → 159.2 RT = 11.9 minutes)

**Figure 18.**
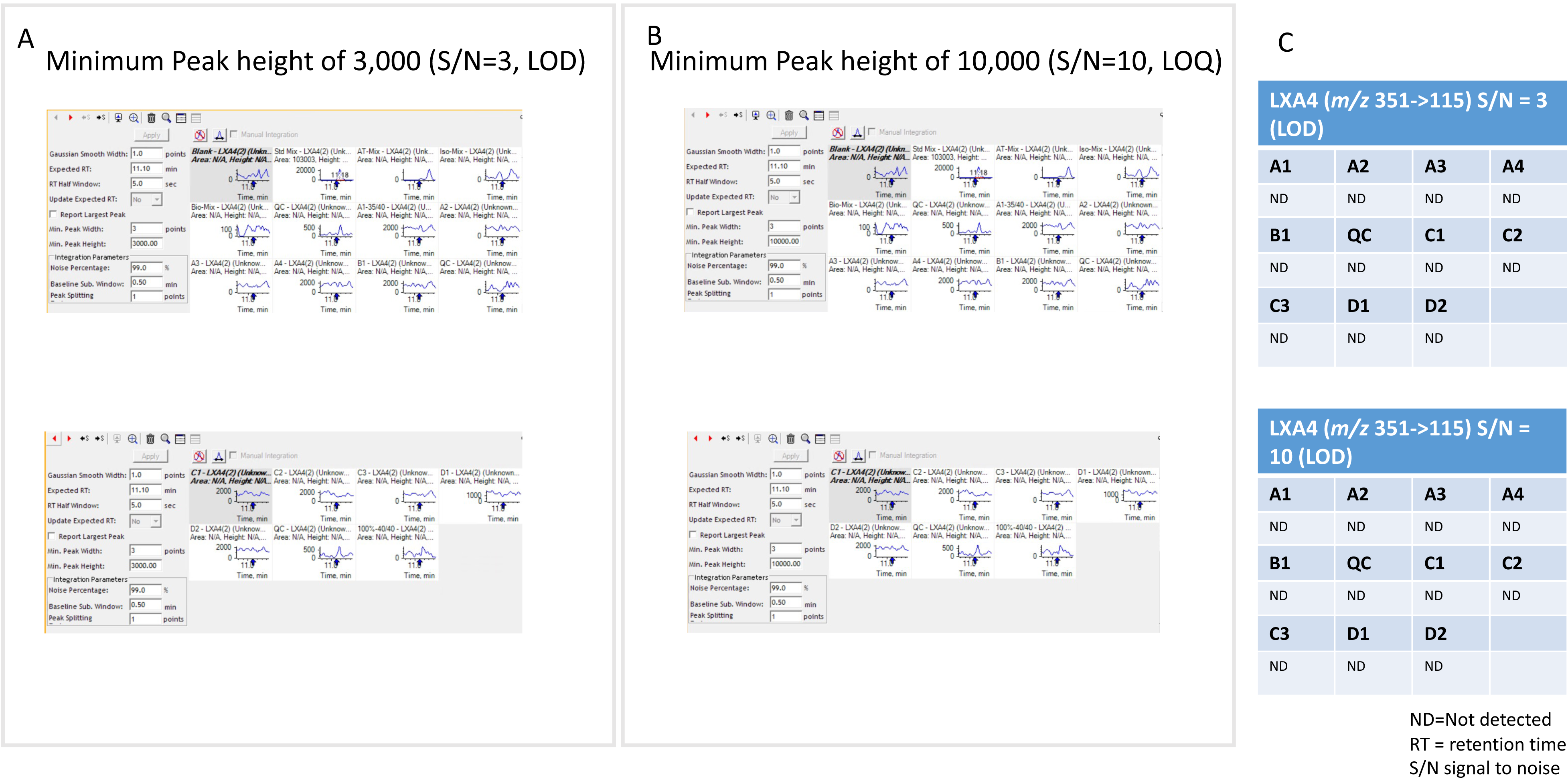
LXA_4_ (*m/z* 351.1 → 115.1 RT = 11.1 minutes)

**Figure 19.**
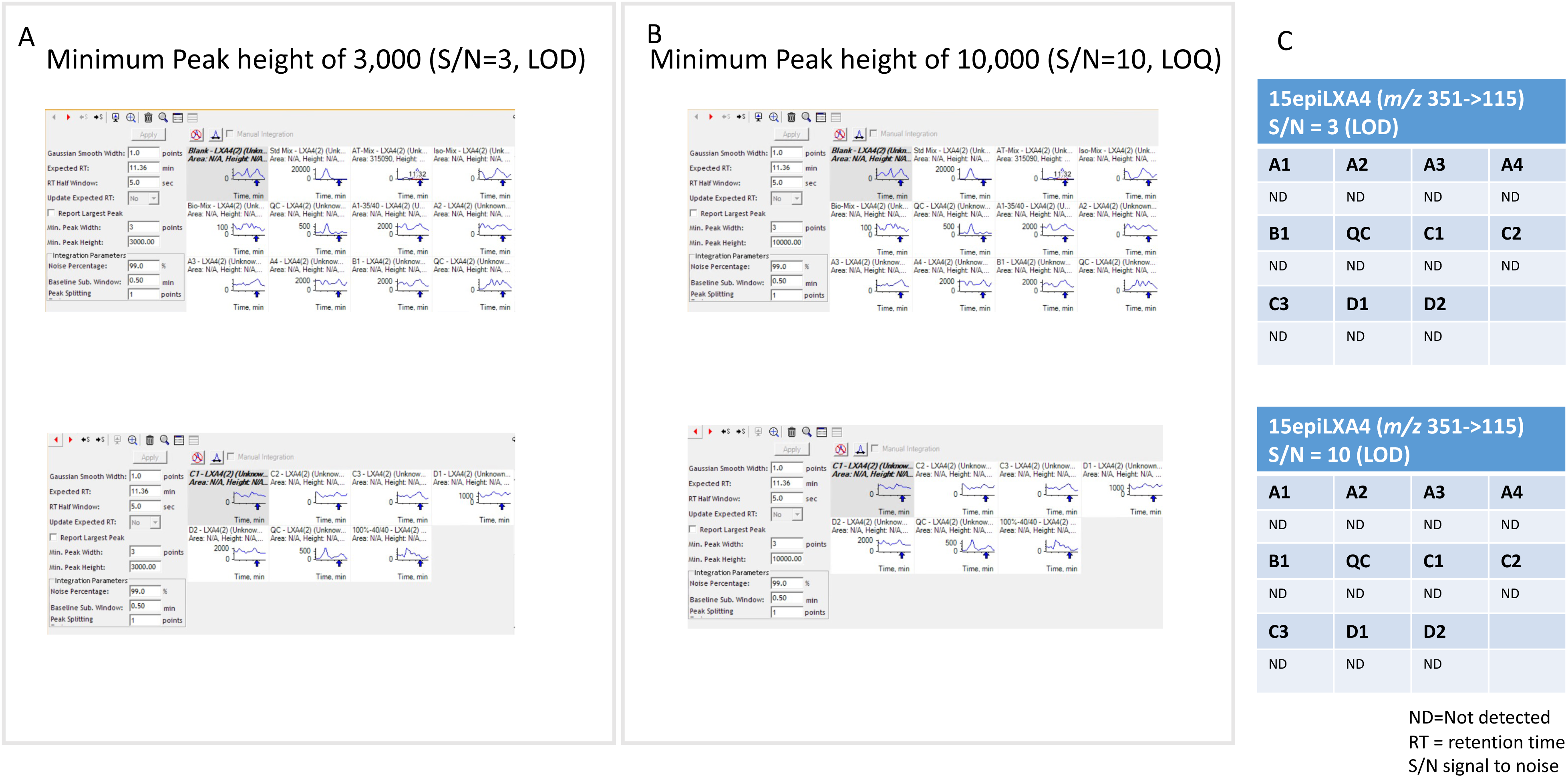
15-epi-LXA_4_ (*m/z* 351.1 → 115.1 RT = 11.36 minutes)

In each data panel the labelling of the subset of data files provided is as follows:

1. Reagent Blank
2. Sets of mixed standard compounds:
  - Std Mix, AT-Mix, Iso-mix, Bio-mix, QC
3. Lipids are labelled as follows:
  - <u>SPMs:</u> RvD1; RvD2; RvD3; RvD4; RvD5; RvD6; 17-RvD1; 17-RvD3; PD1; 10S, 17S-diHDHA; MaR1; 7S, 14S-diHDHA; LXB4; RvE3, 5S,15S-diHETE; RvE1; RvE2; LxA4; 15-epi-LxA4
  - <u>Prostaglandins:</u> PGE_2_, PGD_2_, PGF_2α_ and TxB_2_
  - <u>Leukotriene:</u> LTB_4_
4. Biological samples are labelled as follows:
  - A1,2,3,4 (Corresponding to four 48h time points)
  - B1 (Day14)
  - C1,2,3 (Corresponding to three Day 17 time points)
  - D1,2 (D1 = 8 hours and D2 = 14 hours)

Re-analysis of the data of the samples did not yield detectable peaks (SNR>3) for the docosahexaenoic acid derived resolvins RvD1; RvD2; RvD3; RvD4; RvD6; 17-RvD1; 17-RvD3. Equally, no peaks were detected for the protectins or maresins: PD1; MaR1, 7S, 14S-diHDHA, or the arachidonic acid derived LXA4, LXB4 or 15-epi-LXA4. No peaks were detectable for the eicosapentaenoic acid derived RvE1, RvE2 or RvE3. Peaks for RvD5, and 10S,17S-diHDHA were putatively detected, but did not reach the cut-off SNR 10 to permit quantitation. In contrast, the prostaglandins PGE_2_, PGD_2_ and PGF_2α_ were robustly detected in all or an appropriate subset of samples. TXB_2_ was detected but with poor peak shape. The arachidonic acid derived lipoxin 5S,15S-diHETE was detected with SNR>10.

In terms of applying the unweighted standard curve supplied across the full range, as described in the publication, the relative mean error (RME) of accuracy for the 0.78, 1.56 and 3.12 pg points did not achieve the acceptable limits of <20%. The 1.56 and 3.12 pg points could be brought within 20% limits only if 5 points in the range 1.56-12.5 pg were plotted. Regarding the samples and using the shorter range, amounts in two of the Day 17 samples were in the range allowing quantitation and the rest of the quantities were in the range of the standard curve where RME exceeded 20%.

In summary, the supplementary figure contains peaks for all lipid mediators, but we have not found quantifiable peaks in blister fluid.

## SUMMARY

We(4, 5) showed in rodents that there is a novel phase of prolonged immune activity following resolution of the original inflammatory response that shapes long-term tissue immunity and preserves tissue integrity. We discovered that these events are mediated, in part, by mononuclear phagocyte-derived PGs(5).

The aim of the manuscript in question(1) was to translate these finding to our human model of dermal inflammation. There, we found that once inflammation in response to intradermal UV-KEc had resolved (as defined by clearance of heat, redness, swelling and pain and well as tissue immune cells and inflammatory mediators), there was the infiltration of macrophages with robust prostanoid biosynthesis. As in our rodent studies, blocking post-resolution prostanoids using naproxen revealed a role for these lipids in shaping tissue immunity.

As part of the lipidomic analysis, which was carried out by Professor Dalli, profiles of SPMs were also reported. Reanalysis of these data revealed that the mediators in Figs 4A-D were detected and lay within the range of concentrations anticipated to be quantified robustly, with a caveat that the peaks of TXB_2_ were of poor chromatographic shape. However, peaks were not detected for many SPMs in blister fluid, examined in accordance with harmonised ICH guidelines(2, 3), and results in our conclusion that representation of the presence of SPMs, originally presented in the Supplementary file and the quantities of LXB4, RvD5 and RvE3 in Figs 4E, F and H of this report of human inflammatory exudates traversing three phase of inflammatory onset, resolution and post-resolution, must be ignored. The values presented for quantities of 5,15S-diHETE in Fig 4G are within the calibration curves subject to errors greater than 20% but were detected with SNR>10 and thus must be interpreted with caution.

We present these data for broader academic consideration.

This does not detract for the role of other lipids in post-resolution biology.

**Figure 20.**
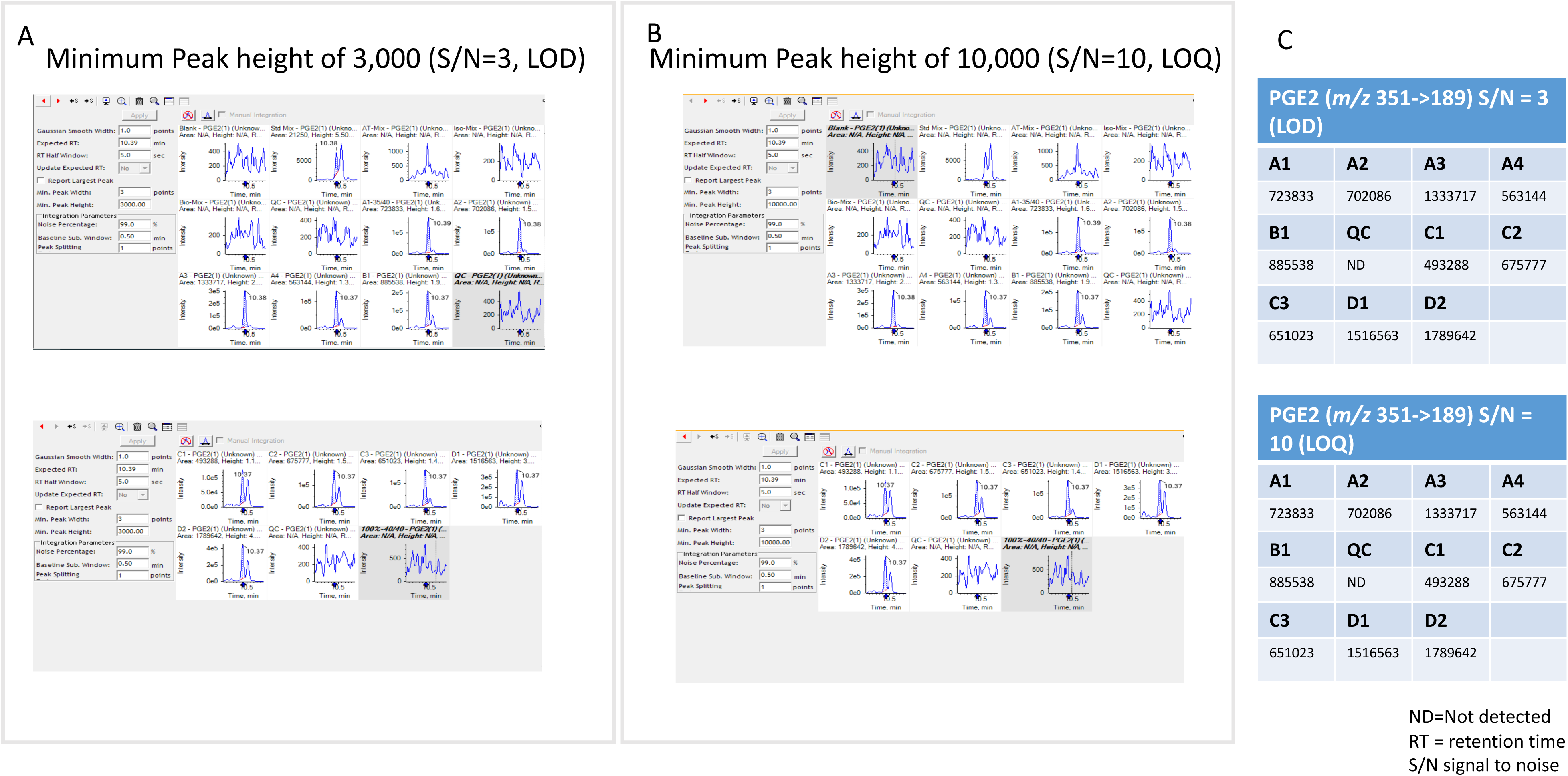
PGE_2_ (*m/z* 351.3 → 189.1 RT = 10.39 minutes)

**Figure 21.**
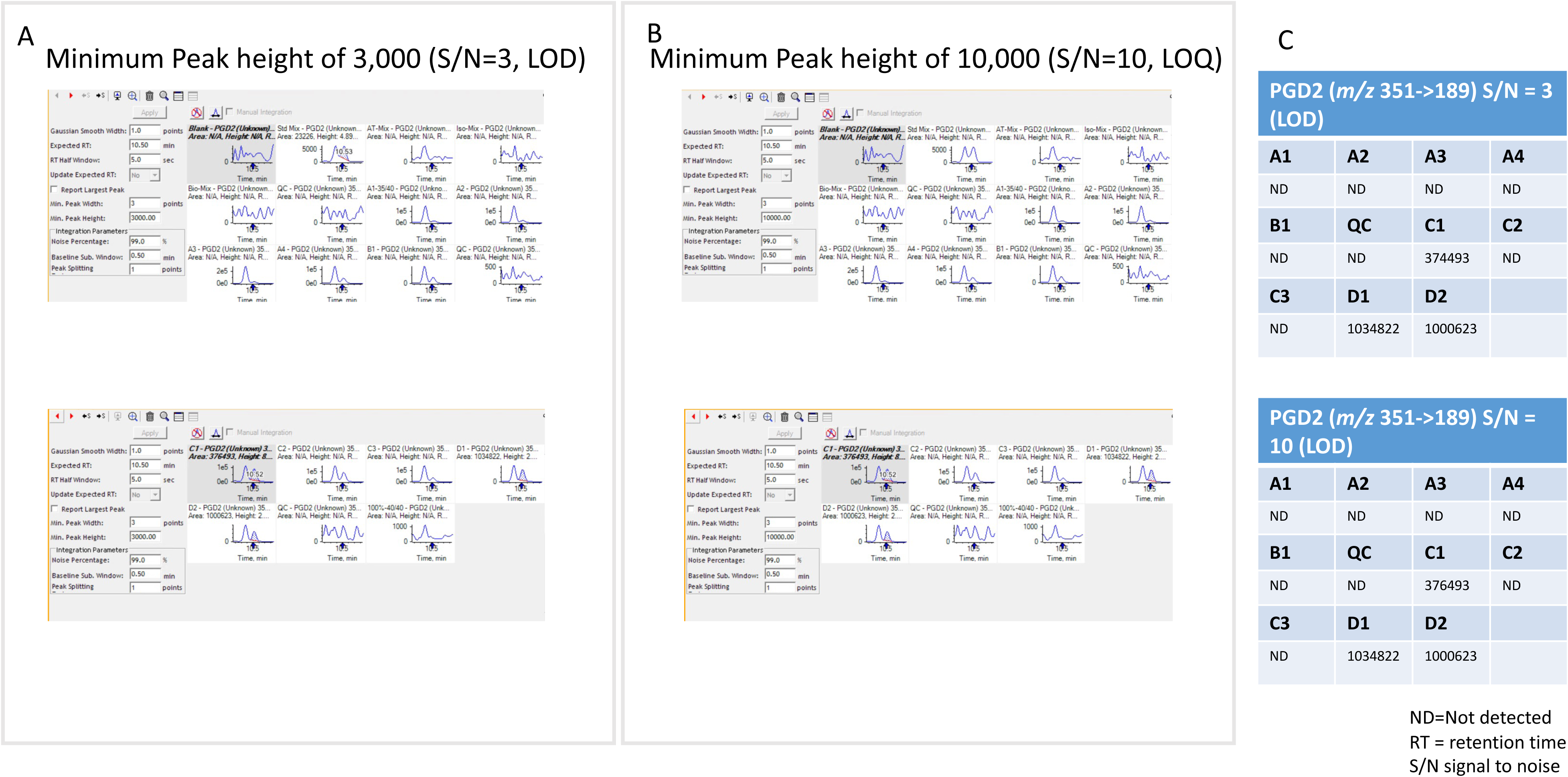
PGD_2_ (*m/z* 351.1 → 189.1 RT = 10.5 minutes)

**Figure 22.**
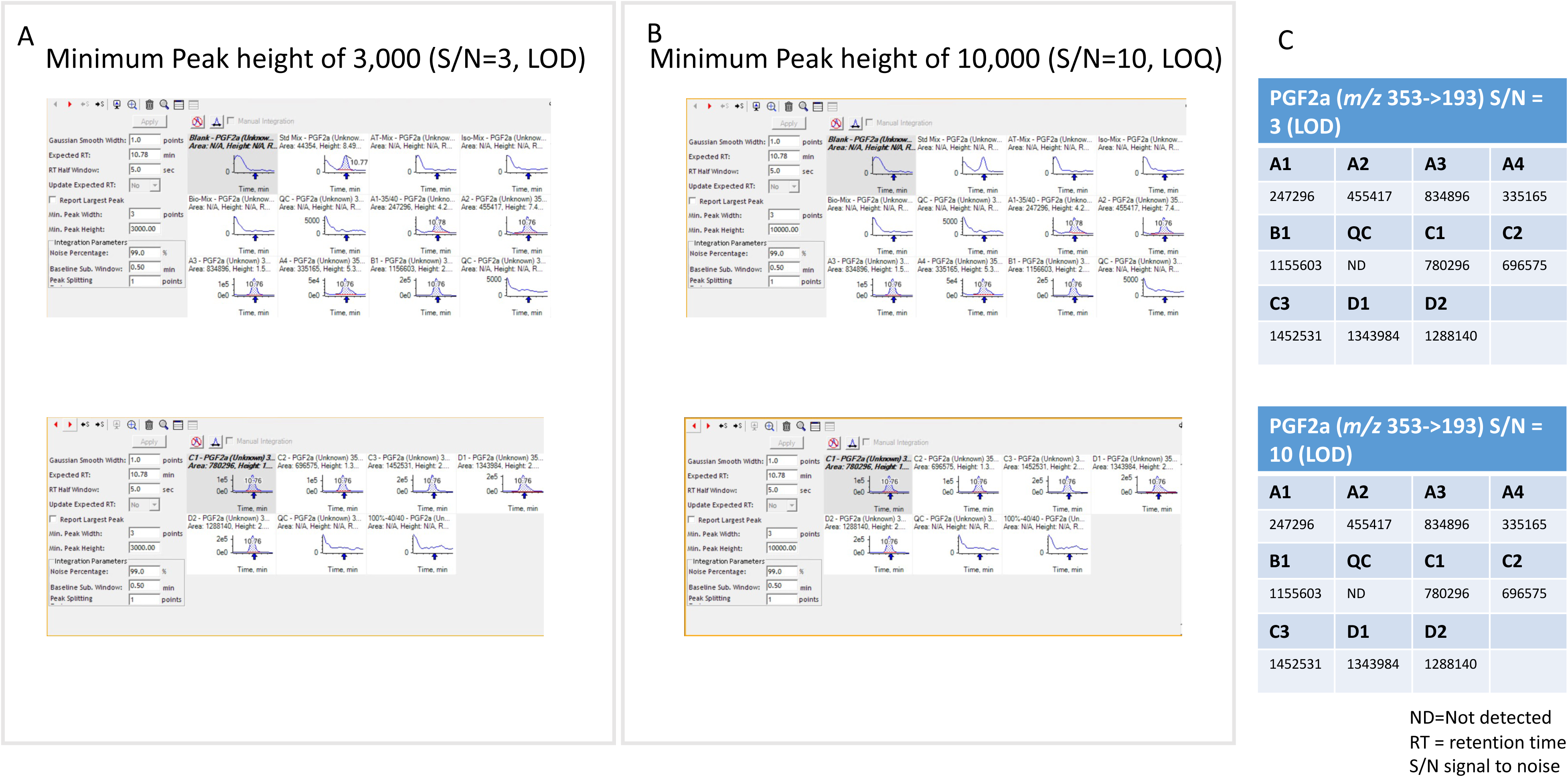
PGF2α (*m/z* 353.3 → 193.1 RT = 10.78 minutes)

**Figure 23.**
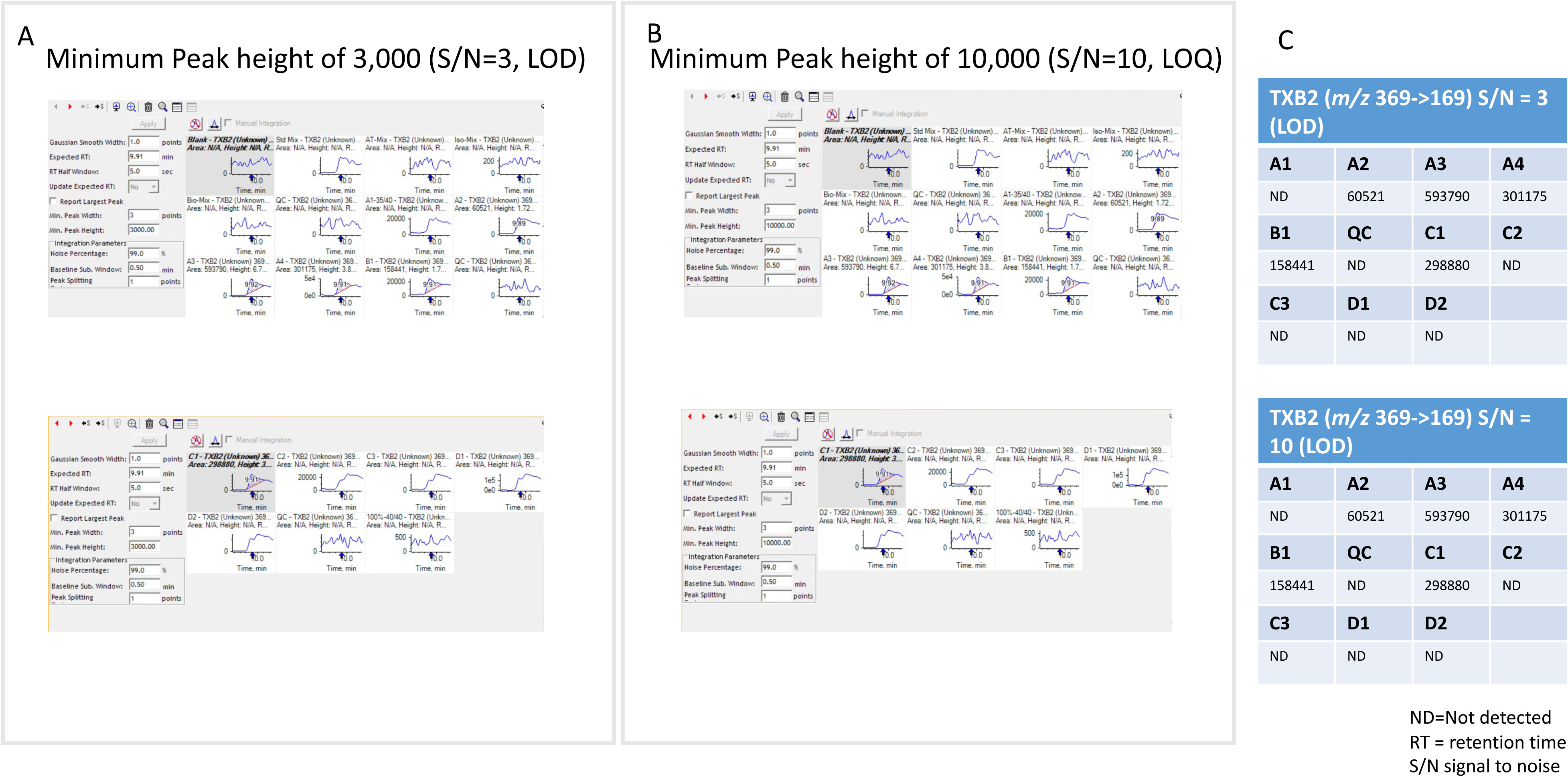
TXB_2_ (*m/z* 369.3 → 169.1 RT = 9.91 minutes)

**Figure 24.**
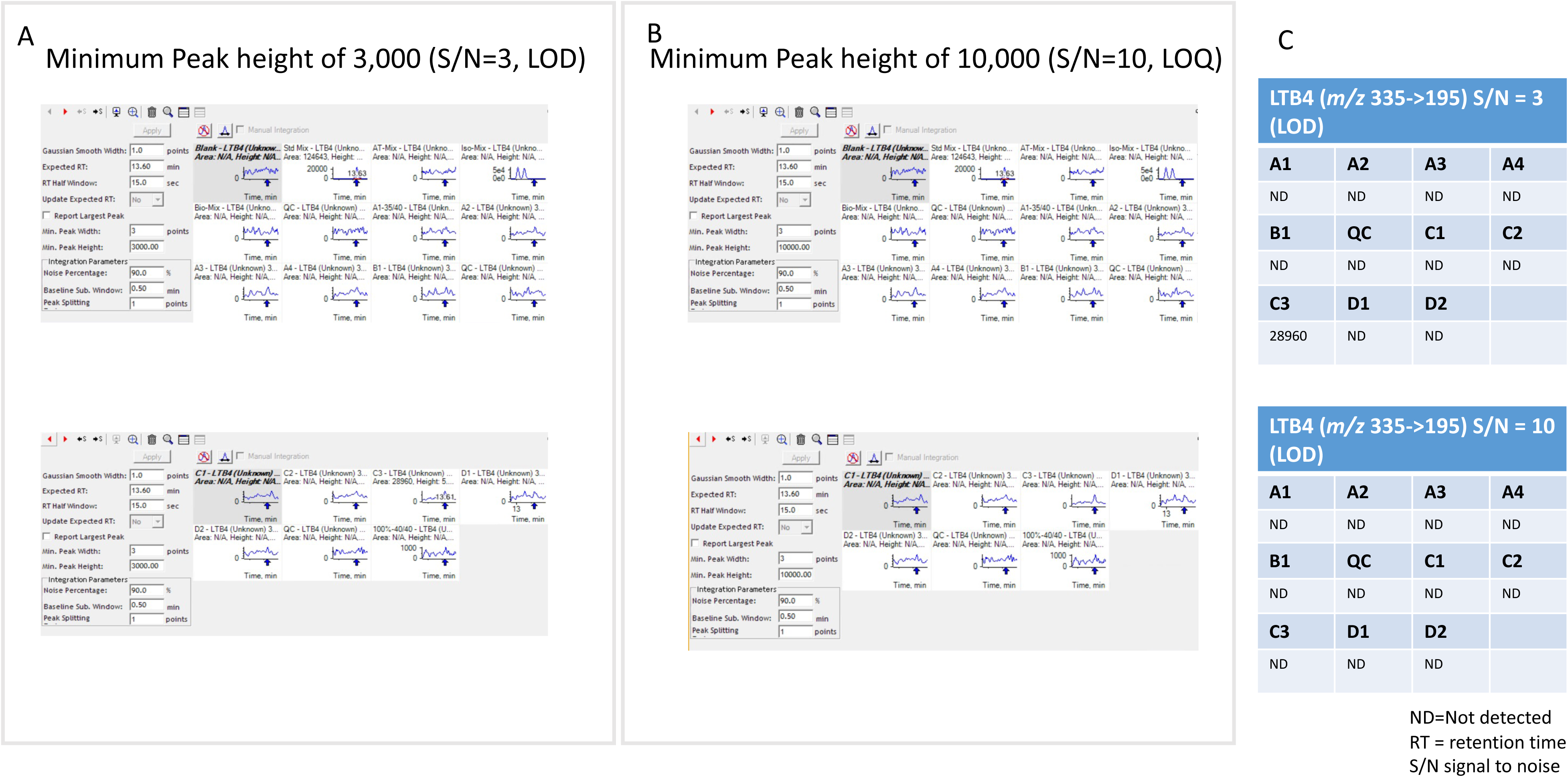
LTB_4_ (*m/z* 335.3 → 195.1 RT = 13.6 minutes)

